# An Introductory Implementation of Breast Cancer Detection from Mammograms and Pixel Intensity with Efficient-Net Other Neural Nets

**DOI:** 10.1101/2024.05.04.592536

**Authors:** Sheekar Banerjee, Humayun Kabir

## Abstract

In the world of civilized medical scientific progression, cancer has become a very serious threat for the natural survival of human beings where breast cancer stays to be the second most dangerous type. Mostly women are embracing very pathetic death because of the delayed detection of the cancer cell in the certain period of their life. Machine Learning mechanism can definitely help at the stage of medical imaging which can escalate the diagnosis of the cancer cells at a very early age of its biological formation and development. We focused upon the deep learning approach to classify the normal and abnormal breast according to the medical imaging from the MIAS dataset of Mammograms and Pixel Intensity. The Convolution Neural Network (CNN) alongside ResNet, AmoebaNet and EfficientNet have been used for the detection with 330 mammograms in which 194 images are normal and 136 are having the identification of abnormal breasts. The accuracy of the entire experimental results was carrying the torch of potential legacy of deep learning in the medical imaging arena. The research is ongoing for the further development and optimization of CNN, AmoebaNet-C and EfficientNet architecture for the Pixel Intensity with higher accuracy, proper segmentation and masking. Source code of this research is available here: https://github.com/ac005sheekar/Breast-Cancer-Detection-with-Pixel-Intensity/

## I. Introduction

One type of illness known as cancer is one in which the majority of the cells produce a lump or mass known as a tumor. When it comes to breast cancer, the symptom is still unknown because there is no early pain. Therefore, if early screening is conducted, early detection is extremely likely. However, not all bumps are malignant. 21 histological subtypes of breast cancer occur, and over 80% of these cells are invasive [1]. Thousands of new cases of invasive breast cancer cells are eventually diagnosed each year in a number of countries. Among all cancer kinds, breast cancer has a greater than 91% chance of being fully cured. Early-stage breast cancer does not cause pain [2]. The estimation of who will survive for a given period of time after the diagnosis reflects the survival rate of patients. Women and men who have the medical history of having diagnosed with breast cancer previously, are proven prone to have the same diagnosis later on. For the diagnosis, the term Mammography seems to be very important in the long run [3]. Mammography is a procedure which is one kind of low dose x-ray by which we can get the visual representation of the breast’s internal structure. The different signal processing techniques like microwave imaging, ultrasound imaging, wavelet transform which shows the time frequency representation of a signal using small waveform called as wavelet and curve-let [**?**]. It helps the breast cancer detection with its low frequency imagery where its degree of localization varies with scale and produce images on different scale [5]. Deep learning is one kind of machine learning technique in which a computer generated model performs the task of classification directly learning from sound, visual data or texts. The models are trained over a large number of datasets and there are so many layers in the CNN architectures [**?**]. Deep learning is used to detect cancer cells automatically with medical imaging and proficient calibration of pixel intensity. The task of training a deep convolution network seems to be very difficult for the start because it requires a huge size of data for training. Deep learning is also being used in the field of bioinformatics and molecular imaging [7].

## II. Related Works

The application of CNN can be noticed in the field of medical imaging since 1990 when the hands on activity was initiated in digital mammography. For CNN, the transferability is one of the most important aspects, embedded in pretrained CNN [8]. In the very recent research works, we can see that transfer learning in the field of medical imaging is categorized in two major classes. First one is to use the pre-trained networks for feature extraction from a specific layer of the network [9]. Those features are used to train the new pattern classifier. The second one is the class where the rest of the pre-trained network is used as same expect the fully connected layers that are replaced with a new logistic layer [10]. In the past, there were many different types of techniques with classifiers have been proposed like wavelet transforms, amalgamation of cosine transform, support vector machine (SVM) and descriptive CNN. Those were utilized for the feature extraction for this dataset. Different kinds of comparisons have been done using dataset extraction with different classifiers [11]. The vector machine approach, the SVM classifier was used which features classification was done for two classes as normal and abnormal [12]. The patches with mammogram were used to produce augmented dataset where the enhancement of contrast was applied on the dataset. One of the methods of them used the discrete wavelet transform which decomposes the enhanced mammogram into four sub-bands [13]. On the other side, the discrete curvelet transform was also a technique where classification was done using SVM layer and last layer to train CNN [14] [15].

Within the wavelet transform domain, a time domain signal is subjected to low-pass and high-pass filters, which separate the low- and high-frequency components of the signal to produce a scaled and shifted version of the original wavelet [16]. The curvelet displays the multi-dimensional characteristics of the specific experimentation and detects the fine ridges with an extremely accurate orientation.

Fuzzy logic is clearly present in the specific field of medical imaging. It demonstrates characteristics of the human thought process, including logic reasoning and hypothesis. Even in cases where insufficient mathematical representation is made, it can nevertheless influence the situational outcome. The development of a model for a fuzzy system is particularly challenging since it requires accurate simulation and adjustment. An combination of intelligent systems, neuro fuzzy logic facilitates logical reasoning [17].

When combined with the evolutionary method, C-means clustering produces superior segmentation efficiency for region extraction and masking [18]. Using the watershed transform for region classification, the diagnosis of breast cancer on three-dimensional breast ultrasonography pictures was carried out to differentiate between the fatty and non-fatty tissues [19]. The application of K-means clustering to thermal imaging of the breast increases the apparent benefit of early cancer cell detection. Here, the breast thermograph’s color analysis is used to separate out the hot region [20]. the recommendation to remove the tumor area based on the ultrasound measurements of the breasts. Multiple characteristics were retrieved from the segmented tumor region, and a regression classifier was constructed for statistical analysis. The segmentation was done to differentiate the tumor from the background tissues.

Another method that was applied to the mammography pictures to improve pixel intensity and cancel noise was dyadic wavelet transform. Using DDSM and MIAS datasets for classification and localization, the self-transfer learning proposal for object localization was also reported.

## III. Methodology

The Mammograms MIAS dataset was used in this study. In image 1, a dataset glitch is shown. To attain the maximum accuracy rate, CNN needs a very large size or amount of data for training. The largest dataset that was readily available online was used for training and testing since there was a dearth of larger datasets. The majority of the images we’ve used are free and open-source. For this process, 330 pictures in total were used. The graphics’ exact dimensions were 224 x 224. About 194 of the total photos were classified as normal, and the remaining 136 as abnormal. The asymmetry symptom, where the breasts have more dense mass, was visible in the abnormal photos. The second form of anomaly that explained the aberrant arrangement of breast tissues was architectural distortion. Another type of calcification involves tiny calcium deposits in the breasts. There were 25 pictures like this. There were 25 photos total. Circumscribed masses are defined as irregularly shaped masses in the breasts. There were 25 pictures of hypothesized masses—masses with ill-defined margins—in the collection. The dataset was meticulously divided into 69.39% for training and 30.61% for CNN’s automatic testing in order to complete the training procedure. While the ultrasonic images of the cancer cell were indexed with warm color, the mammography images were in grayscale.

Out of 330 photos, 69.39% of the dataset from Mammograms MIAS was used for training. The stochastic gradient descent moment was employed for the training process, and the base learning rate, max epochs, and mini batch parameters were tuned to produce the best results. In our study, as Figure 2 illustrates, we trained the CNN starting from the very beginning of our application. The network layers of a CNN function as a detection filter to identify whether particular picture patterns are present. Large, easily understood characteristics are detected by the CNN’s first layer. The following layer picks up extremely little traits, and the final layer is able to describe every categorization that the preceding levels were able to identify. As seen in the CNN architectural formation figure, DCNN has seven layers with weights. Convolutional layers make up the first four layers, whereas linked layers make up the final three. Grayscale images are the DCNN’s inputs.

**Fig. 1.**
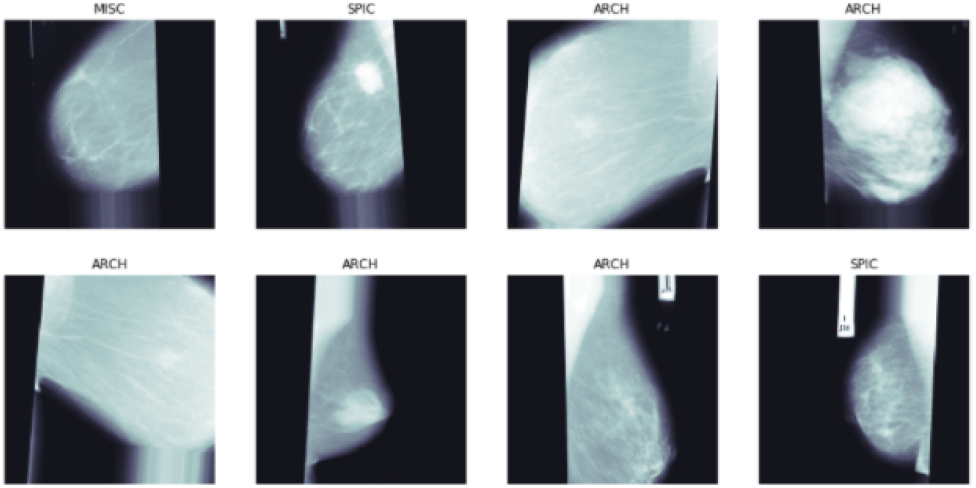
Sample images from the MIAS mammogram dataset.

**Fig. 2.**
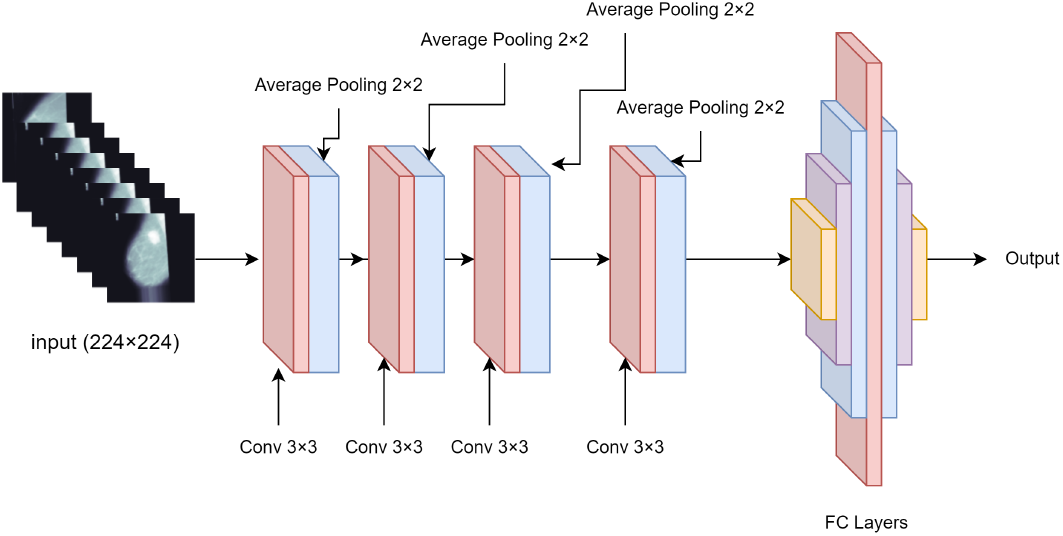
CNN architecture for the proposed research work.

A dot product of weights to the local region connected to the input volume is computed by each neuron. All the borders of the input layer include padding of size (3×3, 2×2, 1×1) and 4, 20, and 80 number of filters of size (2×2, 3×3, 5×5). The [3 × 3] filter sizes specify filters with a height of three and a width of three. The height and width of the input are swept by each filter. In order to save computation and improve resilience, two pooling layers are implemented. Using pooling layers and a 2x2 pixel filter size, each local region receives the maximum value from the four inputs. The learning rate determines how the weights change throughout each epoch in the final layer of the CNN-based classifier. In this case, the weight changes throughout each epoch are defined by the higher learning rate. Notably, we employed a learning rate of 0.01.

The initial input data had dimensions of 224 by 224. The abnormal and normal classes are divided into two groups for the purpose of doing the training. Here, the training set consisted of 94 and 135 photographs, respectively, while the remaining images were used for testing. We attempted to divide each training and testing dataset both automatically and manually by a predetermined ratio, using varying filter sizes, and diverse outcomes were obtained from the process. Our suggested approach is highly effective and has the potential to pave the way for deep learning in the field of cancer cell detection in tomography and mammography. In an effort to improve accuracy outcomes through better CNN architecture optimization, research is still being conducted.

The pre-processed dataset involved resizing the 224x224 input photos during the course of the lengthy processing. The principle component analysis method, or PCA noise filtering, was used to filter out noise from the data. Figure 3 illustrates how the ROI ensemblers determined the pixel intensity from the cell imaging. Seven additional sub-classes were created from the cell dataset in order to assess the accuracy of visual determination within CNN. The pre-processed dataset was used both with and without splitting for training and testing. The figures also demonstrate how excellent the accuracy results were. It was intentional to carry out the segmentation operation by providing raw photos as the input. The morphological closure was completed in this instance. The loss of dilation caused by the morphological closing was crucial to the noise filtering process. Furthermore, the vector transition of the cancer cell in the medical imaging was identified using warm-colored microscopic photography. In this process, the pixel intensity mechanism was started in order to improve the diagnostic outcome of the testing, which was done using training data. The tool or environment utilized for running and implementation is Pycharm community edition. The virtual environment was additionally executed for the purpose of importing integrated CNN libraries, and Python 3 was utilized as the interpreter. When training using an NVIDIA GPU rather than a CPU, a 2GB memory card was utilized as part of the NVIDIA graphics card.

**Fig. 3.**
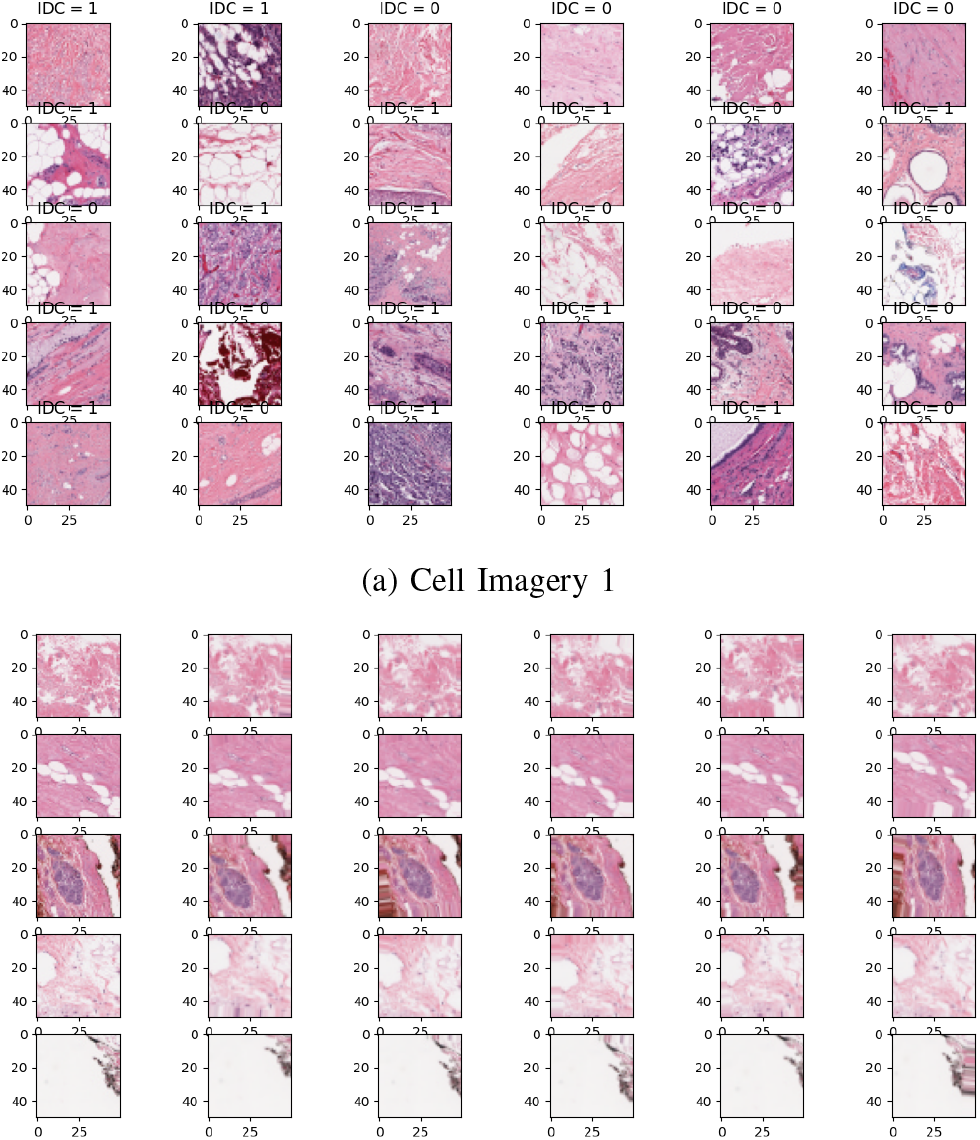
Microscopic imagery sample processing for diagnosis.

## IV. Experiments and Results

The suggested CNN-based breast cancer screening method using pixel intensity features produced extremely positive outcomes. The dataset was subsequently subdivided into 6 additional classes after the aberrant classes were split up into a total of 7 classes. Two classes—normal and abnormal—were used for the training and testing. Each class and subclass underwent unique testing and training. In this case, two distinct scenarios—with and without splitting—were used for the training and assessment. The precision is excellent by all measures.

Every testing iteration demonstrated a very excellent degree of accuracy for the logistic determination of the pixel intensity using the CNN architecture. Ten distinct tests were conducted on the pixel mechanism. The 3D line graphs demonstrate that the 10 iterations yielded an average accuracy of 82%.

In Figure 4, a bar graph and labels show the precision result with accuracy calculation. Figure 5 presents a 3D bar graph and a percentage comparison of CNN accuracies for pre-processed pictures. Figure 6 shows a bar graph representing the pixel intensity from the cell visual data. Figure 7 shows the percentage comparison of CNN accuracies for raw images.

**Fig. 4.**
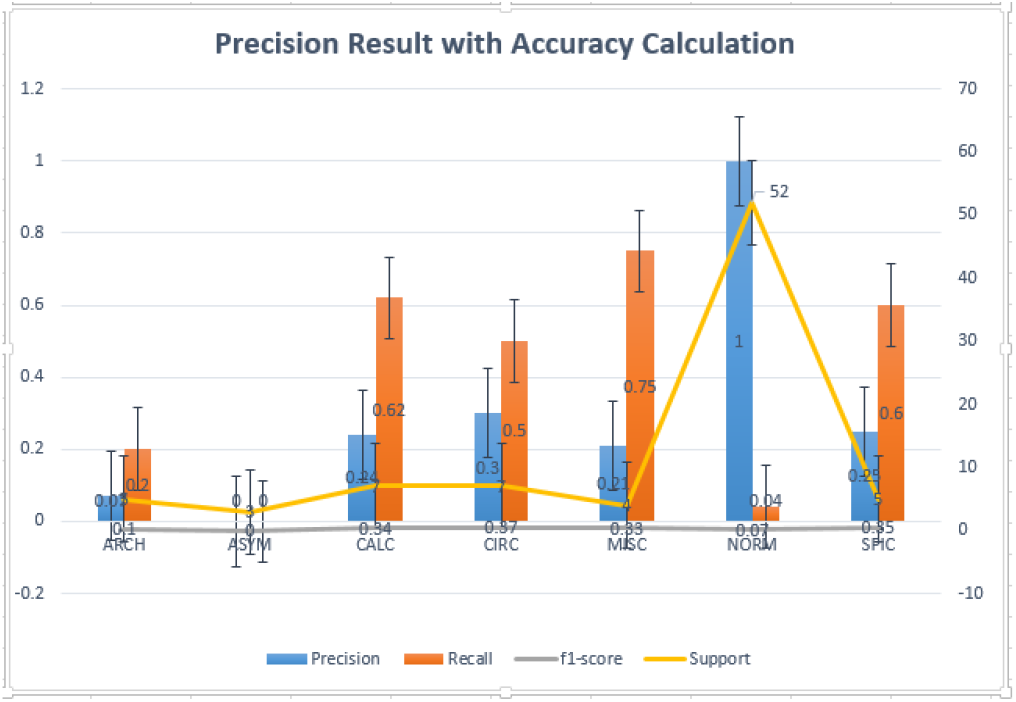
Precision result with accuracy calculation represented with bar graph and labels.

**Fig. 5.**
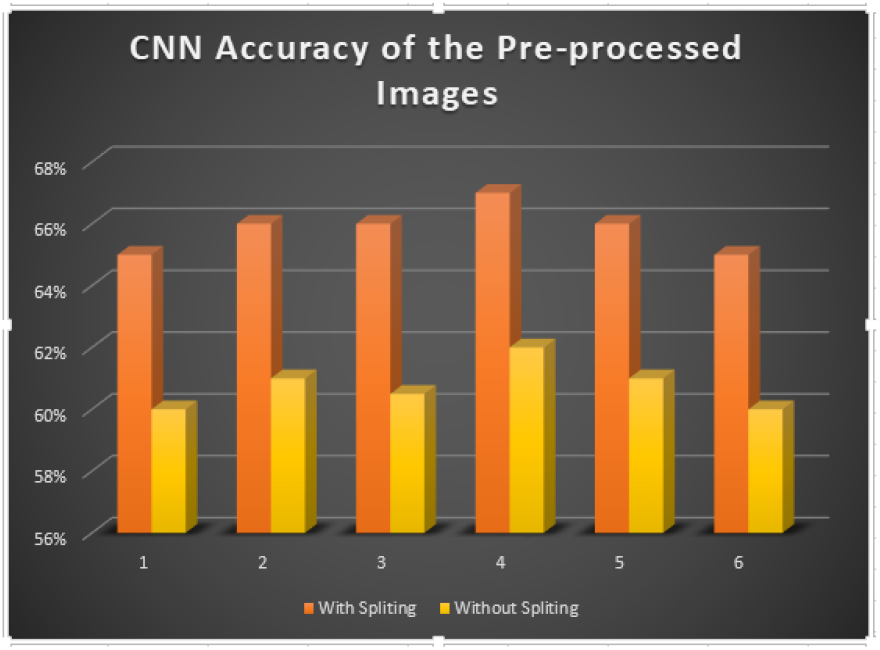
3D bar graph representation of percentage comparison of CNN accuracies for pre-processed images.

**Fig. 6.**
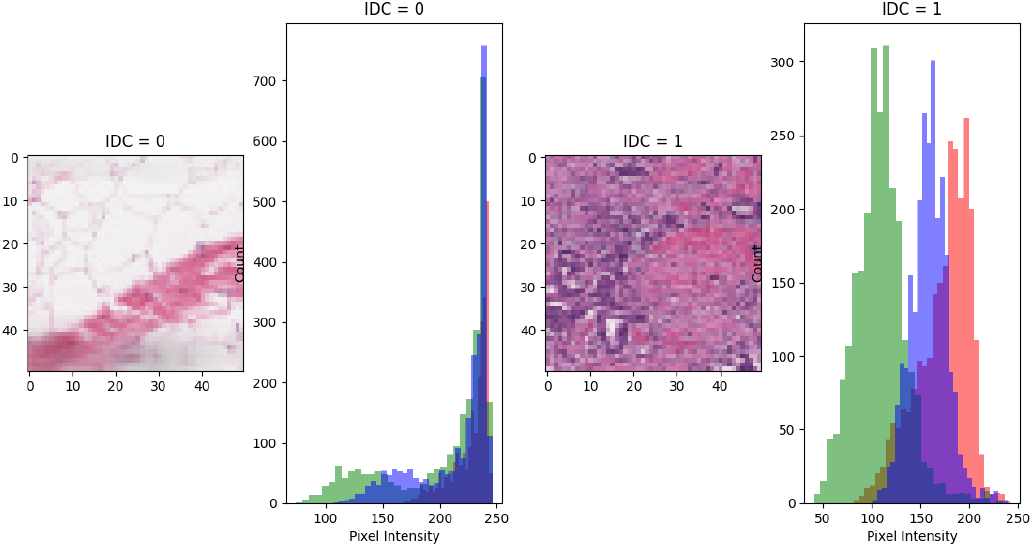
Pixel intensity representation with bar graph from the cell visual data.

**Fig. 7.**
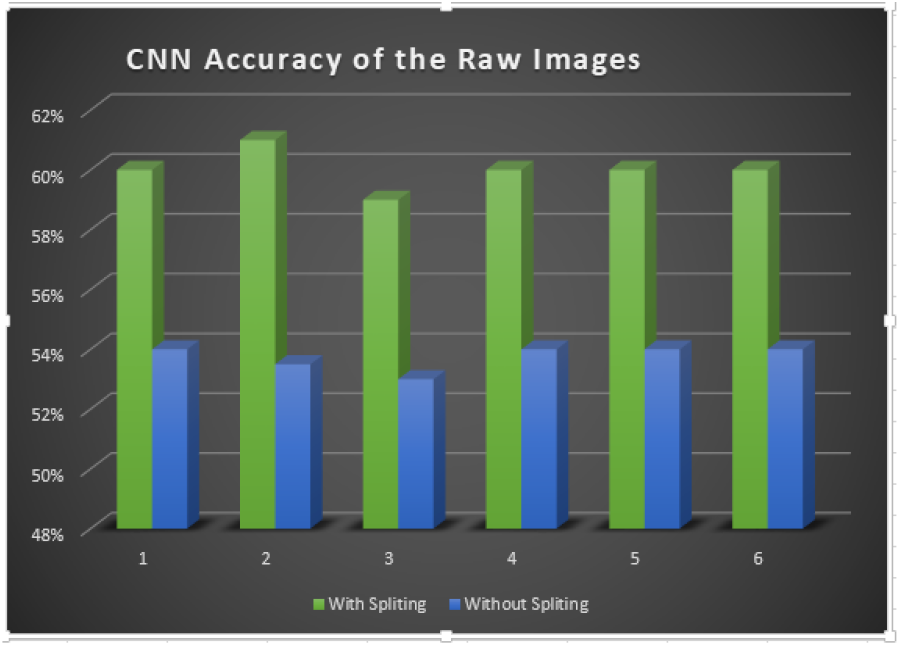
3D bar graph representation of percentage comparison of CNN accuracies for raw images.

Furthermore, Table 3 illustrates the comparison accuracy results between Effective-Net-b7, ResNet101, and Amoeba-Net-C.

**TABLE I.** SPECIFIED REPRESENTATION OF MIAS DATASET WITH TRAIN AND TEST.

| Mammogram Images | Normal | Abnormal |
| --- | --- | --- |
| Training Samples (69.39%) | 135 | 94 |
| Testing Samples (30.61%) | 59 | 42 |

**TABLE II.** TRAINING PARAMETERS FOR CNN.

| Serial No | Parameters | Values |
| --- | --- | --- |
| 1 | Learn rate drop factor | 0.3 |
| 2 | Learn rate drop period | 0.4 |
| 3 | Initial learn rate | 0.01 |
| 4 | Maximum Epochs | 52 |

**TABLE III.**
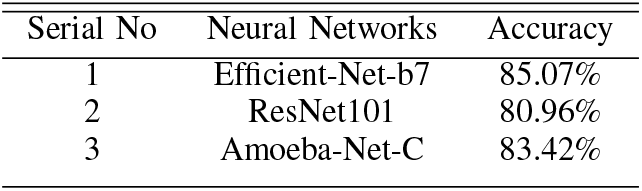
Comparative accuracy results amongst other Neural Networks.

## V. Conclusion and Future works

The entire study project was concentrated on CNN-based categorization using Pixel Intensity and Mammography pictures. The implementation achieved a satisfactory level of success in distinguishing between abnormal and normal mammograms. The mammography MIAS dataset proved to be really helpful in carrying out the study with significant effort and producing results that could be published. The open-source microscopical images datasets have a significant impact on the goal of maintaining pixel intensity stability in the pursuit of improved amalgamation. It is highly likely that this suggested approach will stimulate additional deep learning research in order to make a greater contribution to the field of cancer cell detection, ranging from mammography to any type of tomography, with improved accuracy results and better integration ideas with deep learning in many other bioinformatics and molecular fields.

## Notes

### Competing Interest Statement

The authors have declared no competing interest.

